## Supplementary material for "Restriction of essential amino acids dictates the systemic response to dietary protein dilution": Supp Info

**Supplementary Table 1. Diet formulations from Specialty Feeds.**

| Ingredient | Diet Number | | | | | | | | | | | |
|---|---|---|---|---|---|---|---|---|---|---|---|---|
| | SF17-180 | SF17-175 | SF17-176 | SF17-177 | SF17-179 | SF18-066 | SF18-109 | SF18-110 | SF14-162 | SF17-144 | SF17-115 | SF19-086 |
| | g/Kg | | | | | | | | | | | |
| Sucrose | 333.14 | 237.92 | 337.78 | 340.17 | 339.93 | 340.11 | 340.17 | 340.17 | 100 | 100 | 100 | 340.12 |
| Casein | 190.37 | 47.58 | 48.25 | | | | | | 200 | 34.88 | 159.5 | |
| L Methionine | 2.86 | 0.71 | 3.65 | 4.9 | 5.05 | 4.9 | 1.22 | 1.03 | 3 | 0.78 | 12.48 | 1.22 |
| L Alanine | | | 3.65 | 4.9 | 5.05 | 4.9 | 1.22 | 1.03 | | 3.47 | | 7.52 |
| L Arginine | | | 4.28 | 5.74 | 5.92 | 5.74 | 1.44 | 1.22 | | 4.51 | | 8.82 |
| L Asparagine | | | 5.03 | 6.75 | 6.96 | 6.75 | 1.69 | 1.43 | | 3.74 | | 10.38 |
| L Aspartic Acid | | | 3.65 | 4.9 | 5.05 | 4.9 | 1.22 | 1.03 | | 4 | | 7.52 |
| L Cystine | | | 1.61 | 1.19 | 1.22 | 1.19 | 0.3 | 0.25 | | 0.24 | | 1.82 |
| L Glutamic Acid | | | 14.97 | 20.1 | 20.73 | 20.1 | 5.02 | 4.26 | | 13.4 | | 30.86 |
| L Glutamine | | | 12.32 | 16.55 | 17.07 | 16.55 | 4.14 | 3.51 | | 11.27 | | 25.42 |
| Glycine | | | 2.13 | 2.87 | 2.95 | 2.87 | 0.72 | 0.61 | | 2.22 | | 4.41 |
| L Histidine | | | 3.27 | 4.39 | 4.53 | 4.39 | 1.1 | 0.93 | | 3.22 | | 1.1 |
| L Isoleucine | | | 5.4 | 7.26 | 7.49 | 7.26 | 1.82 | 1.54 | | 7.83 | 1.89 | 1.82 |
| L Leucine | | | 11.32 | 15.2 | 15.68 | 15.2 | 3.8 | 3.22 | | 14.26 | 3.48 | 3.8 |
| L Lysine | | | 9.44 | 12.67 | 13.07 | 12.67 | 3.17 | 2.69 | | 8.96 | | 3.17 |
| L Phenylalanine | | | 6.04 | 8.11 | 8.36 | 8.1 | 2.03 | 1.72 | | 5.83 | | 2.03 |
| L Proline | | | 12.7 | 17.06 | 17.59 | 17.05 | 4.27 | 3.62 | | 12.31 | | 26.2 |
| L Serine | | | 7.17 | 9.63 | 9.94 | 9.63 | 2.41 | 2.04 | | 6.62 | | 14.79 |
| L Threonine | | | 5.15 | 6.92 | 1.73 | 1.73 | 1.73 | 6.29 | | 0.23 | 9.61 | 1.73 |
| L Tryptophan | | | 1.51 | 2.02 | 2.09 | 2.02 | 0.51 | 2.02 | | | 2.03 | 0.51 |
| L Tyrosine | | | 6.54 | 8.79 | 9.06 | 8.78 | 2.2 | 1.87 | | 6.24 | | 13.49 |
| L Valine | | | 6.66 | 8.94 | 9.94 | 8.94 | 2.24 | 1.9 | | 10.44 | 2.54 | 2.24 |
| L Homoserine | | | | | | 5.35 | | | | | | |
| Soybean Oil | 23.8 | 23.79 | 24.13 | 24.3 | 24.28 | 24.29 | 24.29 | 24.29 | 70 | 70 | 70 | 24.29 |
| Lard | 19.04 | 19.03 | 19.3 | 19.44 | 19.42 | 19.43 | 19.44 | 19.44 | | | | 19.44 |
| Cellulose | 47.59 | 47.58 | 48.25 | 48.59 | 48.56 | 48.59 | 48.59 | 48.59 | 50 | 90.65 | 63.24 | 48.59 |
| Wheat Starch | 299.83 | 501.78 | 304 | 306.15 | 305.95 | 306.11 | 423.07 | 423.15 | 404.41 | 403.19 | 403.91 | 306.11 |
| Dextrinised Starch | 33.31 | 71.38 | 33.78 | 34.02 | 33.99 | 34.01 | 43.74 | 43.73 | 132 | 132 | 132 | 34.01 |
| AIN93 Trace Minerals | 1.33 | 1.33 | 1.35 | 1.36 | 1.36 | 1.36 | 1.36 | 1.36 | 1.4 | 1.4 | 1.4 | 1.36 |
| Calcium Carbonate | 5.24 | 5.23 | 5.31 | 5.35 | 5.34 | 5.34 | 5.35 | 5.34 | 13.13 | 13.1 | 13.13 | 5.34 |
| Salt | 2.47 | 2.47 | 2.51 | 2.53 | 2.53 | 2.53 | 2.53 | 2.53 | 2.59 | 2.59 | 2.59 | 2.53 |
| Potassium Dihydrogen Phosphate | | | | | | | | | 6.86 | 10.11 | 6.04 | |
| Sodium Bicrbonate | | | 7.24 | 7.29 | 7.28 | 7.29 | 7.29 | 7.29 | | | | 7.29 |
| Potassium Sulphate | 1.52 | 1.52 | 1.54 | 1.56 | 1.55 | 1.55 | 1.56 | 1.56 | 1.63 | 7.53 | 1.18 | 1.55 |
| Potassium Citrate | 15.71 | 15.71 | 15.92 | 16.04 | 16.03 | 16.03 | 16.04 | 16.03 | 2.48 | 2.48 | 2.48 | 16.03 |
| Dicalcium Phosphate | 12.37 | 12.37 | 12.55 | 12.63 | 12.63 | 12.63 | 12.63 | 12.63 | | | | 12.63 |
| AIN 93 Vitamins | 9.52 | 9.52 | 9.65 | 9.72 | 9.71 | 9.72 | 9.72 | 9.72 | 10 | 10 | 10 | 9.72 |
| Choline Chloride 75% | 1.9 | 1.91 | 1.93 | 1.94 | 1.94 | 1.94 | 1.94 | 1.94 | 2.5 | 2.5 | 2.5 | 1.94 |
| Red Food Colour (124) | | 0.02 | 0.02 | | | | | | | | | |
| Blue Food Colour 10% (133) | | 0.15 | | | | | | | | | | 0.19 |
| Egg Yellow Food Colour (102) | | | | 0.02 | | | | | | | | |
| Green Food Colour (133, 102) | | | | | 0.02 | | | 0.02 | | | | |
| Sunset Yellow Food Colour (110) | | | | | | 0.05 | 0.03 | | | | | |
| Total | 1000 | 1000 | 1000 | 1000 | 1000 | 1000 | 1000 | 1000 | 1000 | 1000 | 1000 | 1000 |

**Supplementary Table 2. Diet formulations from Research Diets.**

| Product # | A14011801 | | A14011802 | | A14011803 | | A14011804 | | A14011806 | |
|---|---|---|---|---|---|---|---|---|---|---|
| | 100% AA | | 25% AA | | 25% EAA, 100% NE | | 100% EAA, 25% NE | | 25% EAA, 154% NE | |
| | 41.7% EAA, 58.3% NE | | 41.7% EAA, 58.3% NE | | 15.2% EAA, 84.8% NE | | 74.1% EAA, 25.9% NE | | 10.4% EAA, 89.6% NE | |
| | gm | kcal | gm | kcal | gm | kcal | gm | kcal | gm | kcal |
| Protein | 16.8 | 17.6 | 4.2 | 4.4 | 11.6 | 12.1 | 9.5 | 9.9 | 16.8 | 17.6 |
| Carbohydrate | 68.7 | 72.1 | 81.3 | 85.3 | 74.0 | 77.6 | 76.1 | 79.8 | 68.7 | 72.1 |
| Fat | 4.4 | 10.3 | 4.4 | 10.3 | 4.4 | 10.3 | 4.4 | 10.3 | 4.4 | 10.3 |
| Total | | 100.0 | | 100.0 | | 100.0 | | 100.0 | | 100.0 |
| kcal/gm | 3.8 | | 3.8 | | 3.8 | | 3.8 | | 3.8 | |
| Ingredient (gm) | | | | | | | | | | |
| L-Histidine-HCl-H2O | 4.52 | 18.08 | 1.13 | 4.5 | 1.13 | 4.5 | 4.52 | 18.08 | 1.13 | 4.5 |
| L-Isoleucine | 7.47 | 29.88 | 1.87 | 7.5 | 1.87 | 7.5 | 7.47 | 29.88 | 1.87 | 7.5 |
| L-Leucine | 15.64 | 62.56 | 3.91 | 15.6 | 3.91 | 15.6 | 15.64 | 62.56 | 3.91 | 15.6 |
| L-Lysine-HCl | 13.04 | 52.16 | 3.26 | 13.0 | 3.26 | 13.0 | 13.04 | 52.16 | 3.26 | 13.0 |
| L-Methionine | 5.04 | 20.16 | 1.26 | 5.0 | 1.26 | 5.0 | 5.04 | 20.16 | 1.26 | 5.0 |
| L-Phenylalanine | 8.34 | 33.36 | 2.09 | 8.4 | 2.09 | 8.4 | 8.34 | 33.36 | 2.09 | 8.4 |
| L-Threonine | 7.12 | 28.48 | 1.78 | 7.1 | 1.78 | 7.1 | 7.12 | 28.48 | 1.78 | 7.1 |
| L-Tryptophan | 2.08 | 8.32 | 0.52 | 2.1 | 0.52 | 2.1 | 2.08 | 8.32 | 0.52 | 2.1 |
| L-Valine | 9.20 | 36.8 | 2.30 | 9.2 | 2.30 | 9.2 | 9.20 | 36.8 | 2.30 | 9.2 |
| L-Alanine | 5.04 | 20.16 | 1.26 | 5.0 | 5.04 | 20.2 | 1.26 | 5.04 | 7.74 | 31.0 |
| L-Arginine | 5.91 | 23.64 | 1.48 | 5.9 | 5.91 | 23.6 | 1.48 | 5.92 | 9.08 | 36.3 |
| L-Asparagine-H2O | 6.95 | 27.8 | 1.74 | 7.0 | 6.95 | 27.8 | 1.74 | 6.96 | 10.68 | 42.7 |
| L-Aspartate | 5.04 | 20.16 | 1.26 | 5.0 | 5.04 | 20.2 | 1.26 | 5.04 | 7.74 | 31.0 |
| L-Cystine | 1.22 | 4.88 | 0.31 | 1.2 | 1.22 | 4.9 | 0.31 | 1.24 | 1.87 | 7.5 |
| L-Glutamine | 17.03 | 68.12 | 4.26 | 17.0 | 17.03 | 68.1 | 4.26 | 17.04 | 26.16 | 104.6 |
| L-Glutamic Acid | 20.68 | 82.72 | 5.17 | 20.7 | 20.68 | 82.7 | 5.17 | 20.68 | 31.76 | 127.0 |
| Glycine | 2.95 | 11.8 | 0.74 | 3.0 | 2.95 | 11.8 | 0.74 | 2.96 | 4.54 | 18.2 |
| L-Proline | 17.55 | 70.2 | 4.39 | 17.6 | 17.55 | 70.2 | 4.39 | 17.56 | 26.96 | 107.8 |
| L-Serine | 9.91 | 39.64 | 2.48 | 9.9 | 9.91 | 39.6 | 2.48 | 9.92 | 15.22 | 60.9 |
| L-Tyrosine | 9.04 | 36.16 | 2.26 | 9.0 | 9.04 | 36.2 | 2.26 | 9.04 | 13.88 | 55.5 |
| Corn Starch | 315 | 1260 | 435.2 | 1741 | 359.2 | 1437 | 381 | 1524 | 315 | 1260 |
| Maltodextrin 10 | 35 | 140 | 45 | 180 | 45 | 180 | 45 | 180 | 35 | 140 |
| Sucrose | 350 | 1400 | 350 | 1400 | 350 | 1400 | 350 | 1400 | 350 | 1400 |
| Cellulose | 50 | 0 | 50 | 0 | 50 | 0 | 50 | 0 | 50 | 0 |
| Soybean Oil | 25 | 225 | 25 | 225 | 25 | 225 | 25 | 225 | 25 | 225 |
| Lard | 20 | 180 | 20 | 180 | 20 | 180 | 20 | 180 | 20 | 180 |
| Sodium Bicarbonate | 7.5 | 0 | 7.5 | 0 | 7.5 | 0 | 7.5 | 0 | 7.5 | 0 |
| S10026 | 10 | 0 | 10 | 0 | 10 | 0 | 10 | 0 | 10 | 0 |
| DiCalcium Phosphate | 13 | 0 | 13 | 0 | 13 | 0 | 13 | 0 | 13 | 0 |
| Calcium Carbonate | 5.5 | 0 | 5.5 | 0 | 5.5 | 0 | 5.5 | 0 | 5.5 | 0 |
| Potassium Citrate, 1 H2O | 16.5 | 0 | 16.5 | 0 | 16.5 | 0 | 16.5 | 0 | 16.5 | 0 |
| Vitamin Mix V10001 | 10 | 40 | 10 | 40 | 10 | 40 | 10 | 40 | 10 | 40 |
| Choline Bitartrate | 2 | 0 | 2 | 0 | 2 | 0 | 2 | 0 | 2 | 0 |
| FD&C Yellow Dye #5 | 0.05 | 0 | 0.025 | 0 | 0 | 0 | 0.025 | 0 | 0 | 0 |
| FD&C Red Dye #40 | 0 | 0 | 0.025 | 0 | 0.05 | 0 | 0 | 0 | 0 | 0 |
| FD&C Blue Dye #1 | 0 | 0 | 0 | 0 | 0 | 0 | 0.025 | 0 | 0.05 | 0 |
| Total | 1038.32 | 3940 | 1038.22 | 3940 | 1038.19 | 3940 | ###### | 3940 | 1038.30 | 3940 |
| Percent EAA (w/w) | 41.7 | | 41.7 | | 15.2 | | 74.1 | | 10.4 | |
| Percent NEAA (w/w) | 58.3 | | 58.3 | | 84.8 | | 25.9 | | 89.6 | |

| Product # | A14011808 | | A16120601 | | A16120602 | | A16120503 | | A17020901 | |
|---|---|---|---|---|---|---|---|---|---|---|
| | 205% EAA, 25% NE | | | | | | | | 100% Lys | |
| | 85.4% EAA, 14.6% NE | | 19.3% EAA, 80.7% NE | | 17.5% EAA, 82.5% NE | | 23.5% EAA, 76.5% NE | | 15.5% EAA, 84.5% NE | |
| | gm | kcal | gm | kcal | gm | kcal | gm | kcal | gm | kcal |
| Protein | 16.8 | 17.6 | 16.8 | 17.6 | 16.8 | 17.6 | 16.8 | 17.6 | 16.9 | 17.7 |
| Carbohydrate | 68.7 | 72.1 | 68.7 | 72.1 | 68.7 | 72.1 | 68.7 | 72.1 | 68.7 | 72.0 |
| Fat | 4.4 | 10.3 | 4.4 | 10.3 | 4.4 | 10.3 | 4.4 | 10.3 | 4.4 | 10.3 |
| Total | | 100.0 | | 100.0 | | 100.0 | | 100.0 | | 100.0 |
| kcal/gm | 3.8 | | 3.8 | | 3.8 | | 3.8 | | 3.8 | |
| Ingredient (gm) | | | | | | | | | | |
| L-Histidine-HCl-H2O | 9.25 | 37.0 | 1.02 | 4.1 | 4.62 | 18.1 | 0.96 | 3.8 | 1.07 | 4.3 |
| L-Isoleucine | 15.31 | 61.2 | 1.68 | 6.7 | 1.72 | 6.9 | 7.47 | 29.9 | 1.77 | 7.1 |
| L-Leucine | 32.03 | 128.1 | 3.52 | 14.1 | 3.60 | 14.4 | 15.64 | 62.6 | 3.70 | 14.8 |
| L-Lysine-HCl | 26.70 | 106.8 | 13.04 | 52.2 | 3.00 | 12.0 | 2.78 | 11.1 | 13.04 | 52.2 |
| L-Methionine | 10.32 | 41.3 | 1.14 | 4.6 | 5.04 | 20.2 | 1.08 | 4.3 | 1.19 | 4.8 |
| L-Phenylalanine | 17.08 | 68.3 | 1.88 | 7.5 | 8.34 | 33.4 | 1.78 | 7.1 | 1.98 | 7.9 |
| L-Threonine | 14.59 | 58.4 | 7.12 | 28.5 | 1.64 | 6.6 | 1.52 | 6.1 | 1.69 | 6.8 |
| L-Tryptophan | 4.27 | 17.1 | 2.08 | 8.3 | 0.48 | 1.9 | 0.44 | 1.8 | 0.49 | 2.0 |
| L-Valine | 18.86 | 75.4 | 2.07 | 8.3 | 2.12 | 8.5 | 9.20 | 36.8 | 2.18 | 8.7 |
| L-Alanine | 1.26 | 5.0 | 6.97 | 27.9 | 7.13 | 28.5 | 6.61 | 26.4 | 7.33 | 29.3 |
| L-Arginine | 1.48 | 5.9 | 8.18 | 32.7 | 8.36 | 33.4 | 7.75 | 31.0 | 8.60 | 34.4 |
| L-Asparagine-H2O | 1.74 | 7.0 | 9.62 | 38.5 | 9.83 | 39.3 | 9.12 | 36.5 | 10.11 | 40.4 |
| L-Aspartate | 1.26 | 5.0 | 6.97 | 27.9 | 7.13 | 28.5 | 6.61 | 26.4 | 7.33 | 29.3 |
| L-Cystine | 0.31 | 1.2 | 1.68 | 6.7 | 1.72 | 6.9 | 1.60 | 6.4 | 1.77 | 7.1 |
| L-Glutamine | 4.26 | 17.0 | 23.57 | 94.3 | 24.09 | 96.4 | 22.33 | 89.3 | 24.77 | 99.1 |
| L-Glutamic Acid | 5.17 | 20.7 | 28.61 | 114.4 | 29.24 | 117.0 | 27.11 | 108.4 | 30.07 | 120.3 |
| Glycine | 0.74 | 3.0 | 4.09 | 16.4 | 4.18 | 16.7 | 3.88 | 15.5 | 4.30 | 17.2 |
| L-Proline | 4.39 | 17.6 | 24.29 | 97.2 | 24.82 | 99.3 | 23.02 | 92.1 | 25.52 | 102.1 |
| L-Serine | 2.48 | 9.9 | 13.71 | 54.8 | 14.01 | 56.0 | 12.99 | 52.0 | 14.41 | 57.6 |
| L-Tyrosine | 2.26 | 9.0 | 12.50 | 50.0 | 12.78 | 51.1 | 11.85 | 47.4 | 13.14 | 52.6 |
| Corn Starch | 315 | 1260 | 315 | 1260 | 315 | 1260 | 315 | 1260 | 315 | 1260 |
| Maltodextrin 10 | 35 | 140 | 35 | 140 | 35 | 140 | 35 | 140 | 35 | 140 |
| Sucrose | 350 | 1400 | 350 | 1400 | 350 | 1400 | 350 | 1400 | 350 | 1400 |
| Cellulose | 50 | 0 | 50 | 0 | 50 | 0 | 50 | 0 | 50 | 0 |
| Soybean Oil | 25 | 225 | 25 | 225 | 25 | 225 | 25 | 225 | 25 | 225 |
| Lard | 20 | 180 | 20 | 180 | 20 | 180 | 20 | 180 | 20 | 180 |
| Sodium Bicarbonate | 7.5 | 0 | 7.5 | 0 | 7.5 | 0 | 7.5 | 0 | 7.5 | 0 |
| S10026 | 10 | 0 | 10 | 0 | 10 | 0 | 10 | 0 | 10 | 0 |
| DiCalcium Phosphate | 13 | 0 | 13 | 0 | 13 | 0 | 13 | 0 | 13 | 0 |
| Calcium Carbonate | 5.5 | 0 | 5.5 | 0 | 5.5 | 0 | 5.5 | 0 | 5.5 | 0 |
| Potassium Citrate, 1 H2O | 16.5 | 0 | 16.5 | 0 | 16.5 | 0 | 16.5 | 0 | 16.5 | 0 |
| Vitamin Mix V10001 | 10 | 40 | 10 | 40 | 10 | 40 | 10 | 40 | 10 | 40 |
| Choline Bitartrate | 2 | 0 | 2 | 0 | 2 | 0 | 2 | 0 | 2 | 0 |
| FD&C Yellow Dye #5 | 0 | 0 | 0 | 0 | 0.025 | 0 | 0 | 0 | 0.025 | 0 |
| FD&C Red Dye #40 | 0.025 | 0 | 0.05 | 0 | 0 | 0 | 0.025 | 0 | 0.025 | 0 |
| FD&C Blue Dye #1 | 0.025 | 0 | 0 | 0 | 0.025 | 0 | 0.025 | 0 | 0 | 0 |
| Total | 1038.31 | 3940 | 1038.29 | 3940 | 1038.3 | 3940 | 1038.29 | 3940 | 1034.01 | 3943 |
| Percent EAA (w/w) | 85.4 | | 19.3 | | 17.5 | | 23.5 | | 15.5 | |
| Percent NEAA (w/w) | 14.6 | | 80.7 | | 82.5 | | 76.5 | | 84.5 | |

| Product # | A17020902 | | A17020903 | | A17041301 | | A17041302 | | A17041303 | |
|---|---|---|---|---|---|---|---|---|---|---|
| | 100% Thr | | 100% Trp | | Low Lys | | Low Thr | | Low Trp | |
| | 13.2% EAA, 86.8% NE | | 11.2% EAA, 88.8% NE | | 38.1% EAA, 61.9% NE | | 39.8% EAA, 60.2% NE | | 41.2% EAA, 58.8% NE | |
| | gm | kcal | gm | kcal | gm | kcal | gm | kcal | gm | kcal |
| Protein | 16.8 | 17.7 | 16.8 | 17.6 | 16.8 | 17.6 | 16.8 | 17.6 | 16.8 | 17.6 |
| Carbohydrate | 68.7 | 72.1 | 68.7 | 72.1 | 68.7 | 72.1 | 68.7 | 72.1 | 68.7 | 72.1 |
| Fat | 4.4 | 10.3 | 4.4 | 10.3 | 4.4 | 10.3 | 4.4 | 10.3 | 4.4 | 10.3 |
| Total | | 100.0 | | 100.0 | | 100.0 | | 100.0 | | 100.0 |
| kcal/gm | 3.8 | | 3.8 | | 3.8 | | 3.8 | | 3.8 | |
| Ingredient (gm) | | | | | | | | | | |
| L-Histidine-HCl-H2O | 1.10 | 4.4 | 1.12 | 4.5 | 4.80 | 19.2 | 4.66 | 18.6 | 4.56 | 18.2 |
| L-Isoleucine | 1.81 | 7.2 | 1.85 | 7.4 | 7.92 | 31.7 | 7.71 | 30.8 | 7.54 | 30.2 |
| L-Leucine | 3.79 | 15.2 | 3.88 | 15.5 | 16.59 | 66.4 | 16.14 | 64.6 | 15.78 | 63.1 |
| L-Lysine-HCl | 3.16 | 12.6 | 3.23 | 12.9 | 3.26 | 13.0 | 13.46 | 53.8 | 13.16 | 52.6 |
| L-Methionine | 1.22 | 4.9 | 1.25 | 5.0 | 5.35 | 21.4 | 5.20 | 20.8 | 5.09 | 20.4 |
| L-Phenylalanine | 2.03 | 8.1 | 2.07 | 8.3 | 8.85 | 35.4 | 8.61 | 34.4 | 8.42 | 33.7 |
| L-Threonine | 7.12 | 28.5 | 1.76 | 7.0 | 7.55 | 30.2 | 1.78 | 7.1 | 7.18 | 28.7 |
| L-Tryptophan | 0.50 | 2.0 | 2.08 | 8.3 | 2.21 | 8.8 | 2.15 | 8.6 | 0.52 | 2.1 |
| L-Valine | 2.23 | 8.9 | 2.28 | 9.1 | 9.76 | 39.0 | 9.49 | 38.0 | 9.28 | 37.1 |
| L-Alanine | 7.51 | 30.0 | 7.67 | 30.7 | 5.35 | 21.4 | 5.20 | 20.8 | 5.09 | 20.4 |
| L-Arginine | 8.81 | 35.2 | 9.00 | 36.0 | 6.27 | 25.1 | 6.10 | 24.4 | 5.96 | 23.8 |
| L-Asparagine-H2O | 10.36 | 41.4 | 10.58 | 42.3 | 7.37 | 29.5 | 7.17 | 28.7 | 7.01 | 28.0 |
| L-Aspartate | 7.51 | 30.0 | 7.67 | 30.7 | 5.35 | 21.4 | 5.20 | 20.8 | 5.09 | 20.4 |
| L-Cystine | 1.81 | 7.2 | 1.85 | 7.4 | 1.29 | 5.2 | 1.26 | 5.0 | 1.23 | 4.9 |
| L-Glutamine | 25.38 | 101.5 | 25.93 | 103.7 | 18.07 | 72.3 | 17.58 | 70.3 | 17.18 | 68.7 |
| L-Glutamic Acid | 30.81 | 123.2 | 31.48 | 125.9 | 21.94 | 87.8 | 21.34 | 85.4 | 20.87 | 83.5 |
| Glycine | 4.40 | 17.6 | 4.50 | 18.0 | 3.13 | 12.5 | 3.04 | 12.2 | 2.98 | 11.9 |
| L-Proline | 26.16 | 104.6 | 26.72 | 106.9 | 18.62 | 74.5 | 18.11 | 72.4 | 17.71 | 70.8 |
| L-Serine | 14.77 | 59.1 | 15.08 | 60.3 | 10.51 | 42.0 | 10.23 | 40.9 | 10.00 | 40.0 |
| L-Tyrosine | 13.47 | 53.9 | 13.76 | 55.0 | 9.59 | 38.4 | 9.33 | 37.3 | 9.12 | 36.5 |
| Corn Starch | 315 | 1260 | 315 | 1260 | 315 | 1260 | 315 | 1260 | 315 | 1260 |
| Maltodextrin 10 | 35 | 140 | 35 | 140 | 35 | 140 | 35 | 140 | 35 | 140 |
| Sucrose | 350 | 1400 | 350 | 1400 | 350 | 1400 | 350 | 1400 | 350 | 1400 |
| Cellulose | 50 | 0 | 50 | 0 | 50 | 0 | 50 | 0 | 50 | 0 |
| Soybean Oil | 25 | 225 | 25 | 225 | 25 | 225 | 25 | 225 | 25 | 225 |
| Lard | 20 | 180 | 20 | 180 | 20 | 180 | 20 | 180 | 20 | 180 |
| Sodium Bicarbonate | 7.5 | 0 | 7.5 | 0 | 7.5 | 0 | 7.5 | 0 | 7.5 | 0 |
| S10026 | 10 | 0 | 10 | 0 | 10 | 0 | 10 | 0 | 10 | 0 |
| DiCalcium Phosphate | 13 | 0 | 13 | 0 | 13 | 0 | 13 | 0 | 13 | 0 |
| Calcium Carbonate | 5.5 | 0 | 5.5 | 0 | 5.5 | 0 | 5.5 | 0 | 5.5 | 0 |
| Potassium Citrate, 1 H2O | 16.5 | 0 | 16.5 | 0 | 16.5 | 0 | 16.5 | 0 | 16.5 | 0 |
| Vitamin Mix V10001 | 10 | 40 | 10 | 40 | 10 | 40 | 10 | 40 | 10 | 40 |
| Choline Bitartrate | 2 | 0 | 2 | 0 | 2 | 0 | 2 | 0 | 2 | 0 |
| FD&C Yellow Dye #5 | 0.025 | 0 | 0 | 0 | 0.025 | 0 | 0.025 | 0 | 0 | 0 |
| FD&C Red Dye #40 | 0 | 0 | 0.025 | 0 | 0.025 | 0 | 0 | 0 | 0.025 | 0 |
| FD&C Blue Dye #1 | 0.025 | 0 | 0.025 | 0 | 0 | 0 | 0.025 | 0 | 0.025 | 0 |
| Total | 1038.60 | 3941 | 1038.31 | 3940 | 1038.33 | 3940 | 1038.31 | 3940 | 1038.32 | 3940 |
| Percent EAA (w/w) | 13.2 | | 11.2 | | 38.1 | | 39.8 | | 41.2 | |
| Percent NEAA (w/w) | 86.8 | | 88.8 | | 61.9 | | 60.2 | | 58.8 | |

**Supplementary Table 3.** Liver amino acid concentrations of mice on a normal control amino acid contain diet and a diet low in Threonine (Low THR) pre-treated with adeno-associated viruses to express yeast THR biosynthetic transcripts (AAV-yTHR1+4)or a negative control (AAV-GFP) in hepatocytes. Data are nmol/g wet weight liver and are mean ± S.E.M. Two-way ANOVA: effect of diet, *p<0.05; effect of AAV, #p<0.05. ME: main effect.

| Amino acid | Normal Amino Acid | | Low THR | | ME |
|---|---|---|---|---|---|
| | AAV-GFP | AAV-yTHR1+4 | AAV-GFP | AAV-yTHR1+4 | |
| **Ala** | 1730 ± 215 | 2507 ± 203 | 1582 ± 197 | 1910 ± 234 | # |
| **Arg** | 4.58 ± 0.45 | 5.58 ± 0.93 | 4.18 ± 0.33 | 4.84 ± 0.56 | |
| **Asp** | 212 ± 24 | 228 ± 26 | 254 ± 14 | 223 ± 22 | |
| **Glu** | 1275 ± 362 | 1577 ± 337 | 1915 ± 269 | 1427 ± 446 | |
| **Gly** | 734 ± 61 | 1103 ± 152 | 920 ± 40 | 806 ± 59 | *,# |
| **His** | 294 ± 36 | 419 ± 54 | 353 ± 33 | 323 ± 24 | |
| **Ile** | 53.9 ± 7.8 | 79.6 ± 13.2 | 67.5 ± 4.0 | 59.1 ± 3.8 | |
| **Leu** | 101 ± 9 | 152 ± 25 | 126 ± 5 | 108 ± 5 | |
| **Lys** | 145 ± 12 | 182 ± 30 | 152 ± 14 | 146 ± 11 | |
| **Met** | 15.8 ± 1.8 | 24.5 ± 3.9 | 15.6 ± 1.7 | 17.4 ± 1.8 | |
| **Phe** | 45.9 ± 2.3 | 66.6 ± 9.2 | 52.2 ± 1.8 | 51.7 ± 3.8 | |
| **Pro** | 72.6 ± 7.2 | 103.0 ± 22.4 | 80.7 ± 5.7 | 81.8 ± 7.4 | |
| **Ser** | 68.2 ± 8.9 | 125.7 ± 20.5# | 112.6 ± 13.7* | 77.8 ± 10.2# | |
| **Thr** | 91.6 ± 13.3 | 154.1 ± 22.8# | 56.2 ± 6.2 | 98.7 ± 11.6 | *,# |
| **Tyr** | 49.4 ± 5.7 | 70.1 ± 6.7# | 56.6 ± 5.8 | 55.3 ± 9.3 | |
| **Val** | 125 ± 16 | 168 ± 28 | 152 ± 8 | 130 ± 15 | |
